# Desiccation-tolerant *Acinetobacter* as a robust chassis for gas-phase bioprocesses

**DOI:** 10.64898/2025.12.24.696357

**Authors:** Shogo Yoshimoto, Hayata Yamada, Shori Inoue, Katsutoshi Hori

## Abstract

**Background:** Gas-phase bioprocesses that immobilize microbial cells on solid carriers enable efficient conversion of volatile or poorly water-soluble substrates, but they also expose cells to fluctuating humidity and transient desiccation that can compromise viability and catalytic performance. In Gram-negative bacteria, especially in pathogens, desiccation is known to impose multifactorial stress, including loss of cellular water and energetics, damage to DNA and proteins, oxidative stress, and disruption of the cell envelope. However, for chassis strains used in or being considered for gas-phase bioprocesses, their desiccation tolerance and responses to desiccation stress remain incompletely defined.

**Results:** Here, we evaluated the viability, energy status, and gas-phase toluene degradation of *Acinetobacter* sp. Tol 5 as a chassis for gas-phase processes after controlled desiccation (8%, 52%, and >95% RH), in comparison with *Acinetobacter baylyi* ADP1, *Pseudomonas putida*, and *Escherichia coli*. To minimize adhesion-related bias in post-desiccation measurements, we used an *ataA*-deficient Tol 5 mutant (Tol 5 Δ*ataA*) for the assays. *Acinetobacter* strains maintained high viability during desiccation for 16 days, whereas *P. putida* and *E. coli* showed significant loss of viability at 8% RH and 52% RH. Intracellular ATP measurements further indicated that desiccation reduced intracellular ATP in all strains, but *E. coli* rapidly exhausted ATP at 8% and 52% RH, whereas *Acinetobacter* strains and *P. putida* retained intracellular ATP. In a gas-phase toluene degradation assay, immobilized Tol 5 retained higher toluene-degrading activity after desiccation than *P. putida* mt-2. Transcriptome profiling revealed a multilayered Tol 5 response involving DNA and RNA maintenance, cell envelope trafficking and remodeling, redox-responsive functions, and broad repression of growth-associated metabolism.

**Conclusions:** Our results highlight *Acinetobacter*, particularly Tol 5, as a promising chassis strain for gas-phase bioprocesses and suggest potential mechanistic targets for stabilizing bacterial cells under low-water-activity conditions.

## Introduction

Microbial bioprocesses are expected to play an important role in bioremediation and sustainable bioproduction (1–3). Gas-related microbial processes have long been developed in the environmental field as biological air-pollution control technologies that remove volatile organic compounds (VOCs), including BTEX (benzene, toluene, ethylbenzene, and xylene) (4, 5). In the context of carbon capture and utilization (CCU), gas fermentation has re-emerged as a practical platform to valorize CO and CO_2_-rich industrial off-gases, and commercialization has progressed to full-scale operation (6–9). Within this broader landscape of gas-based bioprocessing, gas-fed bioprocesses that immobilize bacterial cells at high density on solid carriers or reactor inner surfaces and supply substrates in the gas phase are gaining interest because they enable efficient delivery of poorly soluble or volatile substrates and reduce the impact of gas–liquid mass transfer limitations (10–14). In these systems, cells function as catalysts while being exposed to low-water environments, such as thin aqueous films, microscopic droplets, or near-dry conditions (10, 15). Consequently, their activity is tightly coupled to water activity and osmotic environments that differ substantially from those in bulk liquid culture, and robust environmental stress tolerance is required to maintain viability and metabolic activity after humidity fluctuations and transient desiccation.

*Acinetobacter* sp. Tol 5 is a bacterium that exhibits exceptionally strong adhesiveness and autoagglutination. It adheres firmly to a wide variety of material surfaces via AtaA, a trimeric autotransporter adhesin, enabling robust attachment to diverse abiotic surfaces (16, 17). Tol 5 was originally isolated as a highly efficient aromatic hydrocarbon-degrading strain from a trickle-bed air biofilter (18), and its metabolic pathways have been recently characterized (19, 20). These properties would make Tol 5 a promising chassis for gas-phase bioprocesses using volatile substrates, and a proof-of-concept gas-phase process has been reported (21). However, unlike conventional liquid-phase processes operated with cells suspended in culture media, gas-phase operation presupposes cellular stability under conditions where partial drying and rewetting can occur during operation. Therefore, understanding how chassis strains survive and maintain their biological activities under different humidity levels is essential for designing and operating robust gas-phase bioprocesses.

Bacterial desiccation tolerance has been investigated mainly for pathogens relevant to hospital and food environments. For example, a multilayered survival strategy has been described in *Acinetobacter baumannii* on dry surfaces, involving trehalose-linked osmoprotection, DNA repair systems, proteostasis modules such as chaperones and hydrophilin-like proteins, and viable but non-culturable (VBNC)-like transitions (22–25). Similar but diverse responses to desiccation stress have also been reported for *Staphylococcus* (26), foodborne pathogens that persist in dry matrices, including *Salmonella* (27), *Cronobacter* (28, 29), and *Listeria* (30), and some extremophile bacteria (31, 32). Taken together, desiccation tolerance is thought to rely on a coordinated set of responses that limit macromolecular damage and preserve recovery capacity, including compatible solute-based osmoprotection, antioxidant defenses, DNA repair, protein quality control, and metabolic remodeling under low water activity. However, the composition, timing, and relative contribution of these modules are highly species- and condition-dependent, and remain poorly defined for bioproduction-relevant chassis operating in low-water environments.

In this study, under controlled humidity conditions, we investigated viability and energy status, assessed post-desiccation gas-phase toluene degradation, and characterized transcriptomic responses to desiccation stress.

## Materials and methods

### Bacterial strains and culture conditions

Bacterial strains used in this study are listed in Table 1. *Escherichia coli* was cultivated in Lysogeny broth (LB) medium at 37 °C with shaking at 140 rpm. *Acinetobacter* sp. Tol 5 and its derivatives were cultivated in LB medium at 28 °C with shaking at 115 rpm. *Pseudomonas putida* and *Acinetobacter baylyi* ADP1 were cultivated in LB medium at 30 °C with shaking at 115 rpm.

**Table 1.**
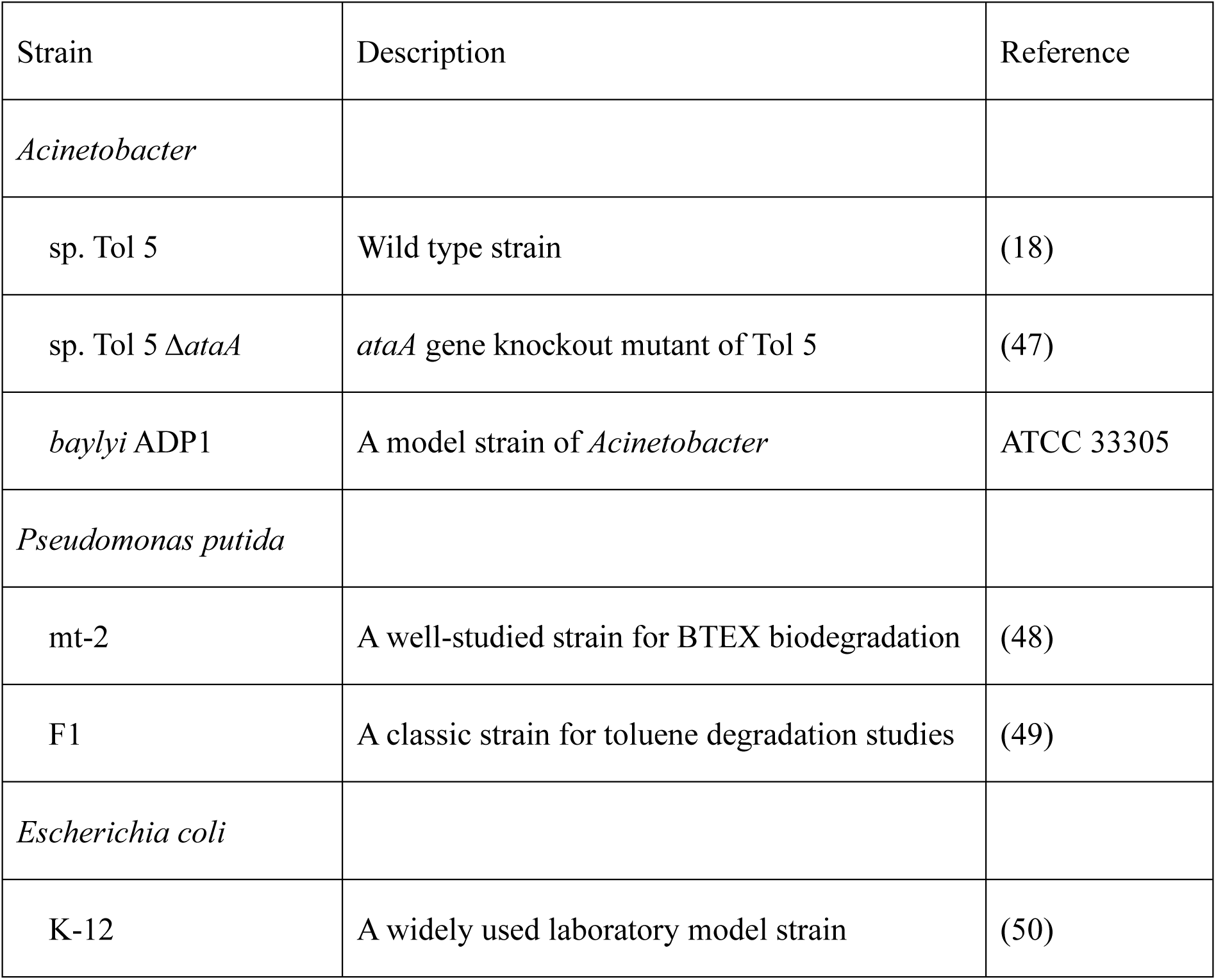
Bacterial strains used in this study.

| Strain | Description | Reference |
| --- | --- | --- |
| *Acinetobacter* |  |  |
| sp. Tol 5 | Wild type strain | (18) |
| sp. Tol 5 Δ*ataA* | *ataA* gene knockout mutant of Tol 5 | (47) |
| *baylyi* ADP1 | A model strain of *Acinetobacter* | ATCC 33305 |
| *Pseudomonas putida* |  |  |
| mt-2 | A well-studied strain for BTEX biodegradation | (48) |
| F1 | A classic strain for toluene degradation studies | (49) |
| *Escherichia coli* |  |  |
| K-12 | A widely used laboratory model strain | (50) |

### Desiccation

An overnight preculture was prepared by cultivating cells in 2 mL of LB medium for 10 h. The culture was then inoculated 1:100 into 20 mL LB medium and incubated for 16 h. Cells were harvested by centrifugation (3,000×g, 10 min) and washed twice with phosphate-buffered saline (PBS) (Nippon Gene, Tokyo, Japan). Washed cells were resuspended in PBS to 1×10^11^ CFU/mL, and 10 µL aliquots were dispensed into a 96-well plate (353072; Corning Inc., Corning, NY, USA). Plates were placed in a 3-L desiccator (017430-150; Sibata Scientific Technology, Soka, Saitama, Japan) and incubated at 28 °C under controlled humidity conditions using the saturated salt solution method (33). Saturated solutions of potassium sulfate (30 g in 8 mL distilled water), magnesium nitrate (25 g in 3 mL distilled water), or lithium bromide (8 g in 1 mL distilled water) were placed at the bottom of the desiccator. The desiccator was sealed using a silicone stopper (6-336-05; AS ONE, Osaka, Japan) equipped with a sensor (SK-L700R-TH-2; Sato Keiryoki Mfg., Tokyo, Japan), and humidity was recorded every hour using a temperature and humidity data logger (SK-L700R-TH; Sato Keiryoki Mfg.). At designated time points, plates were removed from the desiccator, 100 µL of PBS was added to each well, and samples were resuspended by pipetting and then rehydrated by leaving the plate at room temperature for 20 min.

### Colony forming unit (CFU) counting

Serial dilutions were prepared in PBS, and 100 µL of each dilution was spread onto R2A agar plates (396-01611; Shiotani M.S., Amagasaki, Hyogo, Japan). Plates were incubated overnight, colonies were counted, and CFU/mL was calculated taking the dilution factor into account.

### ATP measurement

Cell suspensions (1×10^7^ CFU/mL before desiccation) were mixed 1:1 with BacTiter-Glo Microbial Cell Viability Assay reagent (Promega, Madison, WI, USA) by adding 100 µL of each to a white 96-well plate (353296; Corning). After incubation for 3 min, luminescence was measured using a plate reader (Infinite 200 PRO M Plex; Tecan Group, Männedorf, Switzerland). Luminescence values were normalized to the pre-desiccation signal (set to 100%) to calculate relative ATP levels.

### Toluene degradation assay

Cells were precultured overnight in a 50-mL centrifuge tube containing 4 mL of LB medium supplemented with 2 µL of toluene. The culture was then inoculated (1:100) into 100 mL of LB medium containing 50 µL of toluene in a 500-mL baffled flask. The flask was sealed with a Viton stopper and incubated at 28 °C with shaking at 115 rpm for 16 h. After measuring OD_660_, 5 mL of culture (adjusted to OD_660_ = 2.0) was immobilized onto a glass filter (1825-047; Cytiva, Marlborough, MA, USA) by vacuum filtration. The filter with immobilized cells was incubated in a desiccator under humidity-controlled conditions as described for the desiccation assay. To induce toluene catabolic genes during incubation, 50 µL of toluene was placed inside the desiccator. After 24 h, 600 µL of PBS was added to the filter for rehydration, and the glass filter was transferred to a 50-mL vial and sealed with a Teflon-laminated cap (5-112-7; Maruemu, Osaka, Japan). Toluene (5 µL) was then added using a microsyringe and incubated at 28 °C.

Toluene in the headspace was quantified using a gas chromatograph equipped with a flame ionization detector (GC-FID) (GC-2014; Shimadzu, Kyoto, Japan). An HP-5MS UI capillary column (30 m length, 0.25 mm inner diameter; Agilent, Santa Clara, CA, USA) was used with nitrogen as the carrier gas. Headspace gas (50 µL) was sampled from the vial using a gas-tight syringe (1710N PST-2; Hamilton, Reno, NV, USA) and injected into the GC-FID system. The oven temperature was held at 50 °C for 1 min and then increased to 80 °C at 20 °C/min. The carrier gas total flow rate was 19 mL/min, and the split ratio was set to 5:1.

### RNA sequencing

The desiccated Tol 5 cells in a 24-well plate (92424; TPP Techno Plastic Products, Trasadingen, Switzerland) were resuspended in PBS and immediately collected by centrifugation (10,000×g, 25 °C, 3 min), and the cell pellets were stored at -80 °C. For each desiccation condition, three biological replicates were prepared. Total RNA was extracted using the Cica Geneus RNA Prep Kit (for Tissue) 2 (Kanto Chemical, Tokyo, Japan) according to the manufacturer’s protocol. Ribosomal RNA was removed using the NEBNext rRNA Depletion Kit (Bacteria) (New England Biolabs, Ipswich, MA, USA), and cDNA libraries were generated with the NEBNext Ultra II RNA Library Prep Kit for Illumina (New England Biolabs, Ipswich, MA, USA). The cDNA libraries were sequenced on the Illumina NextSeq550. Raw FASTQ reads were quality-filtered using fastp (version 0.23.2) and mapped to the Tol 5 genome (accession: AP024708 and AP024709) using Bowtie2 (version 2.5.1) with default parameters. Mapped reads were counted using featureCounts (version 2.0.6).

### Differential gene-expression analysis

Raw read counts of CDSs were analyzed in R using the edgeR package (version 3.42.4). Genes with extremely low expression were removed using the filterByExpr function. Normalization was performed using the trimmed mean of M-values (TMM) method implemented in the calcNormFactors function. Differential gene-expression analysis was conducted using the quasi-likelihood pipeline. A design matrix was constructed with the >95% RH condition as the reference and the 8% RH and 52% RH conditions as contrasts. Dispersion estimates were obtained using the estimateDisp function, and model fitting was performed with the glmQLFit function. Differential gene-expression between each condition was assessed using the glmQLFTest function. Genes with FDR < 0.01 and |log2 FC| > 1 were extracted as significantly differentially expressed genes. Gene functions were annotated using multiple databases, including the NCBI database, UniProtKB/Swiss-Prot database, KEGG database, and eggNOG database, as reported previously (19). In addition, genes were categorized by RPS-BLAST searches against the clusters of orthologous groups (COGs) database (34) using COGclassifier (35).

## Results

### Desiccation tolerance under different humidity conditions

To evaluate the impact of desiccation on cell viability, we first quantified colony-forming units (CFU) after desiccation under controlled humidity. Cell suspensions were dispensed into 96-well plates and dried in sealed desiccators whose relative humidity (RH) was controlled by the saturated salt solution method using potassium sulfate (>95% RH), magnesium nitrate (52% ± 1.1%), or lithium bromide (8% ± 0.6%). After desiccation in desiccators, wells were rehydrated by adding PBS and incubating for 20 min, and the recovered cells were plated on agar media for CFU counting. As reference strains, we included *Acinetobacter baylyi* ADP1 as a model *Acinetobacter* strain, *Pseudomonas putida* F1 and mt-2 as Gram-negative bacteria known for aromatic metabolism and biotransformation, and *Escherichia coli* K-12 as a standard laboratory host. Because Tol 5 exhibits strong, nonspecific surface adhesion mediated by AtaA, which could reduce cell detachment and dispersion during rehydration and thereby bias measurements, we used an *ataA*-deficient Tol 5 mutant (Tol 5 Δ*ataA*) for the following assays.

At the lowest humidity (8% RH), *Acinetobacter* strains showed approximately 10^2^-fold and 10^3^-fold higher CFU values than *P. putida* and *E. coli* after 1 day of desiccation (Fig. 1A). As the desiccation period increased, these differences became more pronounced, indicating marked strain-dependent loss of viability under severe desiccation. A similar overall trend was observed at the intermediate humidity condition (52% RH). After 1 day of desiccation, the *Acinetobacter* strains retained CFU values about 10-fold higher than *P. putida* and about 10^4^-fold higher than *E. coli* (Fig. 1B). While the *Acinetobacter* strains maintained CFU levels of roughly 10^8^ CFU/mL under both 8% and 52% RH, *P. putida* declined to approximately 10^2^ CFU/mL at 52% RH after 16 days of desiccation. In particular, *E. coli* dropped below the detection limit after 5 days of desiccation.

**Figure 1.**
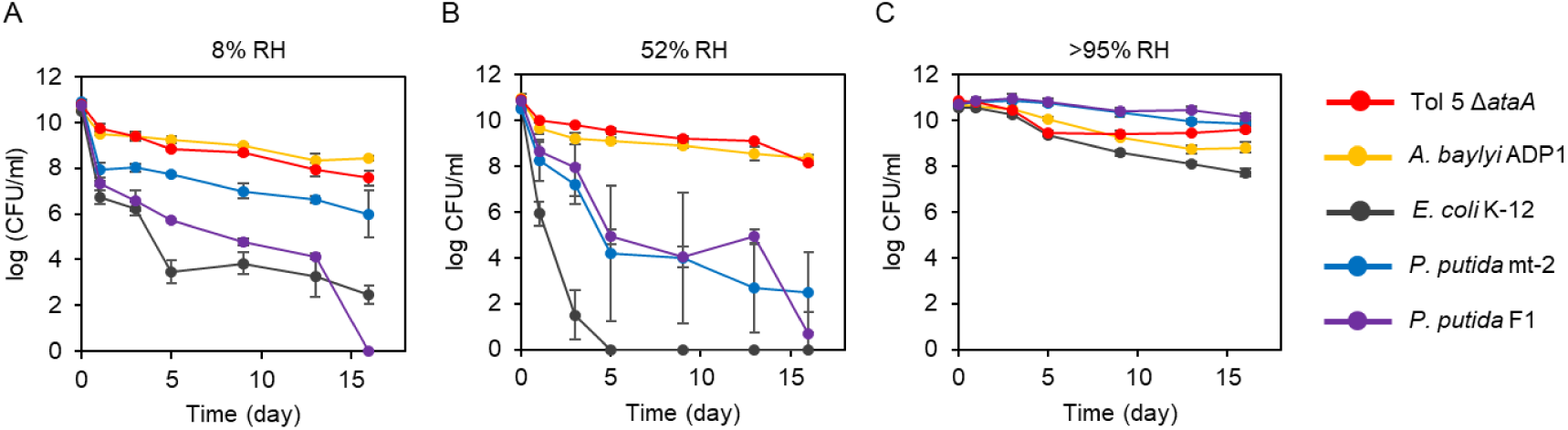
Time-course changes in CFU during desiccation. CFU were quantified after desiccation for the indicated durations. Cells were incubated at (A) 8% RH, (B) 52% RH, or (C) >95% RH. The vertical axis shows CFU, and the horizontal axis shows the days of desiccation. Error bars indicate mean ± standard deviation (biological replicates, n = 3).

In contrast, under the highest humidity condition (>95% RH), all strains maintained CFU values close to those before desiccation after 1 day of desiccation, and high viability was observed even after 16 days of desiccation. The largest decrease was observed for *E. coli*, which still retained 5.5×10^7^ CFU/mL after 16 days of desiccation (Fig. 1C). These results indicate that the >95% RH condition imposed little desiccation stress on the tested bacteria. The observation that the strongest lethal effect occurred at 52% RH is consistent with classical studies reporting that mid-range humidity can be most detrimental to airborne bacteria (36, 37), supporting the notion that distinct physicochemical stress environments arise at different RH levels. Overall, the high viability of the *Acinetobacter* strains across different humidity highlights their strong desiccation tolerance relative to the other tested Gram-negative bacteria.

### ATP levels after desiccation

Next, we measured intracellular ATP levels after desiccation. Because ATP reflects not only cell survival but also the extent to which cells retain metabolic potential after stress exposure, ATP measurements provide a complementary metric to CFU-based assays. After 1 day of desiccation, ATP levels decreased to less than half of the pre-desiccation levels in all strains under all humidity conditions (Fig. 2). Consistent with the significant decrease in CFU at 8% RH and 52% RH, *E. coli* showed an almost complete depletion of ATP under these conditions (Fig. 2A, B).

**Figure 2.**
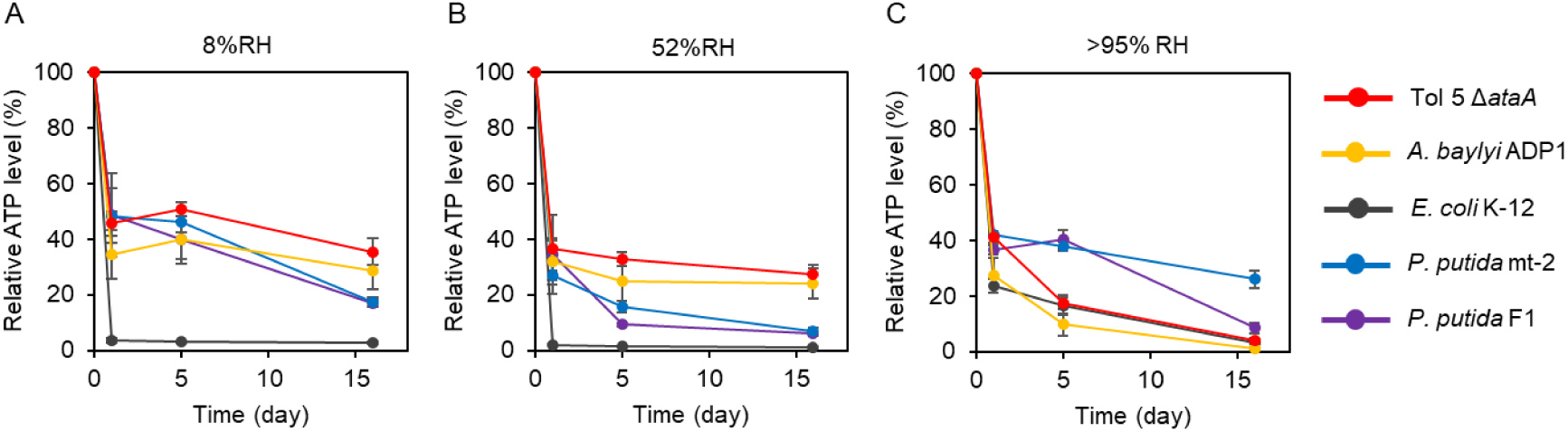
Time-course changes in intracellular ATP levels during desiccation. Intracellular ATP was measured after desiccation. Cells were incubated at (A) 8% RH, (B) 52% RH, or (C) >95% RH before the assay. The vertical axis shows relative ATP levels normalized to the pre-desiccation value (set to 100%), and the horizontal axis shows the days of desiccation. Error bars indicate mean ± standard deviation (biological replicates, n = 3).

Comparing *Acinetobacter* strains and *P. putida*, the *Acinetobacter* strains retained relatively higher ATP levels than *P. putida* after desiccation at 8% RH and 52% RH, whereas at >95% RH *P. putida* maintained higher ATP levels than the *Acinetobacter* strains (Fig. 2C). Together, these results suggest that *Acinetobacter* and *P. putida* may regulate energy metabolism during desiccation rather than undergoing complete ATP exhaustion, although the extent and humidity dependence of ATP retention differ between genera.

### Toluene-degradation activity after desiccation

We assessed the impact of desiccation on gas-phase biocatalysis by measuring the ability of immobilized cells to degrade gaseous toluene as a model substrate. Cells were immobilized on glass fiber filters by vacuum filtration, incubated under the three RH conditions, and then sealed in vials containing toluene. The degradation efficiency was calculated from the decrease in headspace toluene concentration. In this assay, we used the toluene-degrading strains Tol 5 Δ*ataA* and *P. putida* mt-2.

Before desiccation, both strains completely degraded toluene within 21 h (Fig. 3A). After 1 day of desiccation, however, the degradation rate decreased under all RH conditions (Fig. 3B–D). When degradation efficiencies were compared at 48 h, Tol 5 Δ*ataA* degraded toluene almost completely at >95% RH, whereas the degradation efficiencies were 58.4 ± 5.4% at 52% RH and 36.3 ± 5.8% at 8% RH. The delayed initiation of toluene degradation at 8% RH is likely attributable to the time required for recovery from desiccation stress. *P. putida* mt-2 showed nearly complete degradation at >95% RH, but only 47.2 ± 10.2% at 52% RH and 14.1 ± 2.9% at 8% RH. These results indicate that both strains retained measurable catalytic activity after desiccation, with Tol 5 exhibiting slightly higher tolerance to desiccation than mt-2 under the tested conditions.

**Figure 3.**
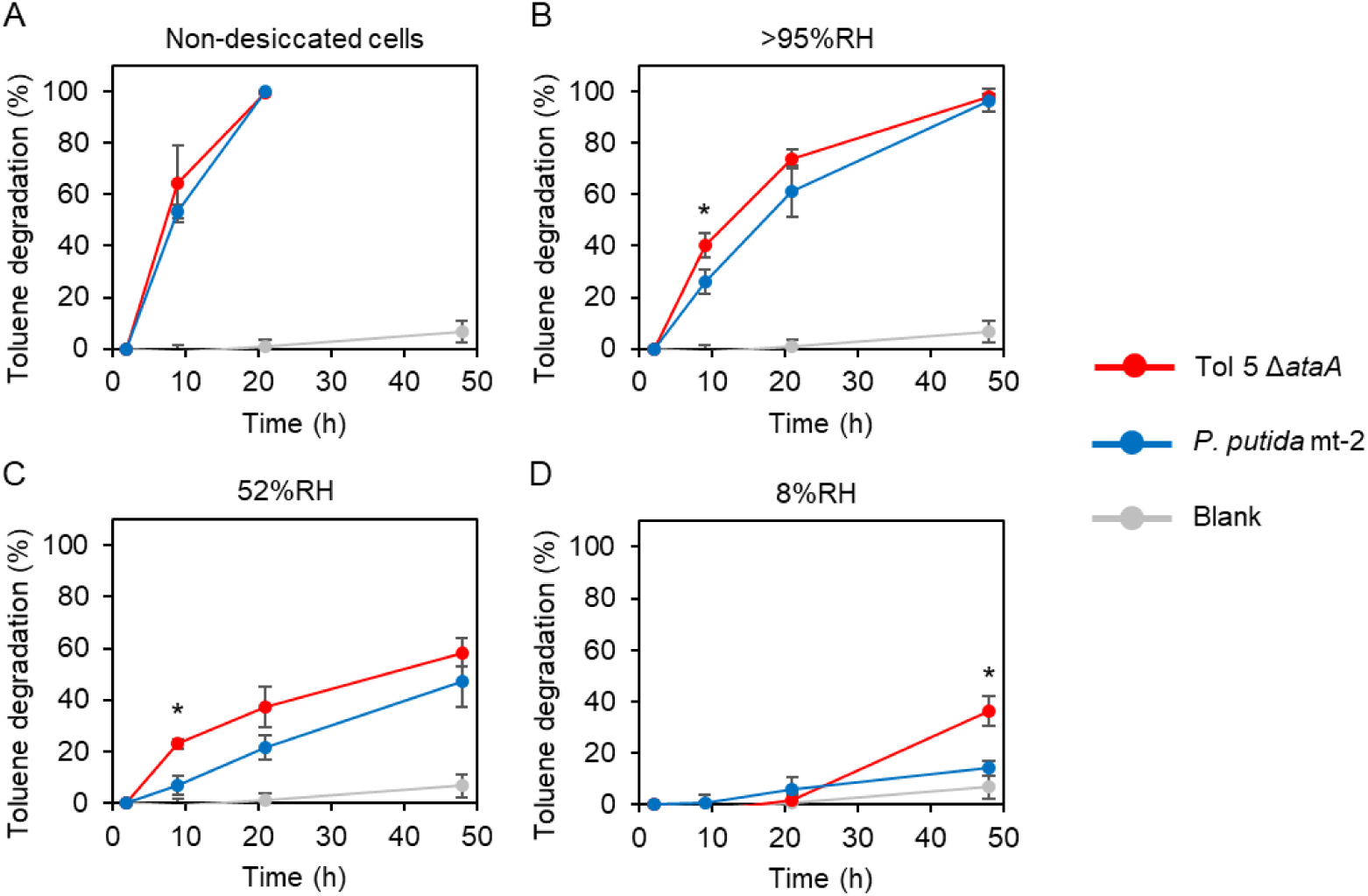
Toluene degradation ability after desiccation. Toluene degradation was monitored by measuring the decrease in toluene after sealing immobilized cells in vials with toluene. (A) Non-desiccated cells were used as a control. Cells were incubated for 1 day at (B) >95% RH, (C) 52% RH, or (D) 8% RH, before the assay. The blank shows the results of incubation without cells. The vertical axis shows the toluene degradation efficiency calculated from the decrease in headspace toluene concentration, normalized to the headspace concentration measured at 2 h after toluene addition. The horizontal axis shows time after toluene addition. Error bars indicate mean ± standard deviation (biological replicates, n = 3). Asterisks represent the statistical significance (*P* < 0.05, Welch’s *t*-test).

### Transcriptomic responses to desiccation stress

We performed whole-genome transcriptome analyses of Tol 5 Δ*ataA* cells after desiccation at >95% RH, 52% RH, and 8% RH, and the expression profiles of all genes were summarized in Table S1. Principal component analysis (PCA) of the transcriptome profiles revealed clear condition-dependent separation, with the >95% RH samples clustering apart from the 52% RH and 8% RH samples. Biological replicates grouped tightly within each condition, indicating good reproducibility of the expression profiles (Fig. 4A), consistent with our observations that desiccation stress was substantially stronger at 52% RH and 8% RH than at >95% RH. We next identified differentially expressed genes (DEGs) at 8% RH and 52% RH based on the thresholds of |log_2_ fold change (FC)| > 1 and false discovery rate (FDR) < 0.01 using the >95% RH condition as the reference. Volcano plots comparing the 8% RH and 52% RH conditions to the >95% RH reference showed the magnitude and significance of differential expression across the genome (Fig. 4B). Under these criteria, 98 genes were significantly altered at 8% RH (52 upregulated and 46 downregulated), whereas 146 genes were significantly altered at 52% RH (61 upregulated and 85 downregulated), with 65 genes (27 upregulated and 38 downregulated) shared between the two conditions. To characterize the functional composition of the DEGs, we classified them based on Clusters of Orthologous Genes (COGs) (34) and summarized the distributions in a pie chart (Fig. 4C). Upregulated genes were enriched for functions related to DNA, RNA, and cell envelopes, whereas downregulated genes were dominated by metabolism-related functions. In addition, DEGs were distributed across diverse functional categories, indicating that Tol 5 exhibits a broad, multilayered response to desiccation stress. The following sections describe these DEGs in detail by functional category.

**Figure 4.**
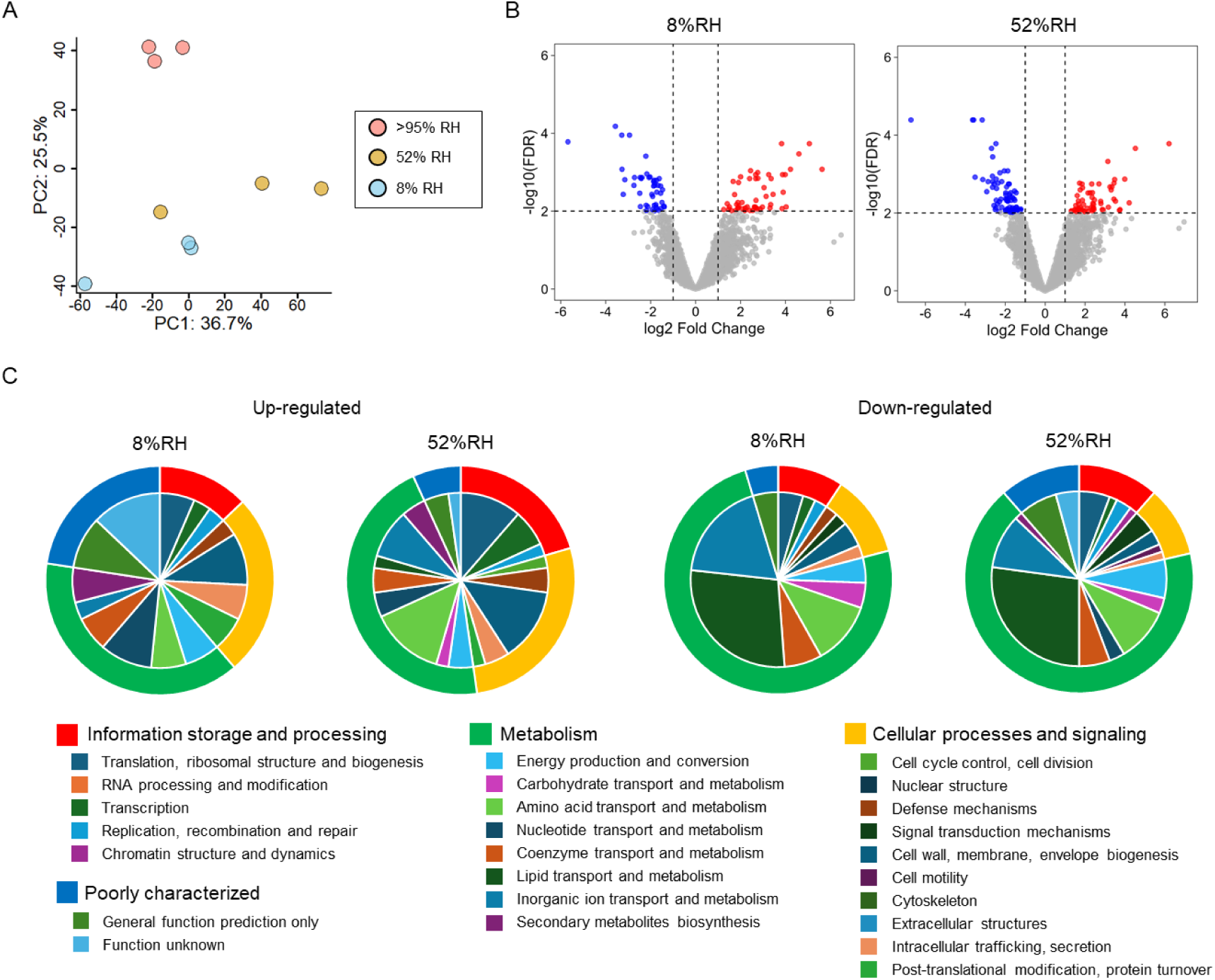
Transcriptomic responses of Tol 5 to desiccation at different relative humidities. (A) PCA of whole-genome transcriptome profiles of Tol 5 after desiccation at >95% RH, 52% RH, and 8% RH. Each point represents an independent biological replicate. The percentage of variance explained by each principal component is indicated on the axes. (B) Volcano plots of differential gene expression at 8% RH (left) and 52% RH (right) relative to the >95% RH reference. Red and blue dots represent significantly upregulated and downregulated genes, respectively (|log_2_FC| > 1, and FDR < 0.01). (C) COG-based functional classification of DEGs identified in (B). Donut charts summarize the functional composition of DEGs for each condition. The outer ring shows the distribution across the four COG supercategories, and the inner ring shows the distribution across the 25 COG functional categories.

DEGs assigned to the “Information storage and processing” supercategory are summarized in Table S2. At 8% RH, the expression of *recA*, which is involved in DNA repair, and *rtcB_2*, which is involved in RNA repair, was markedly upregulated. In addition, transcriptional regulation-related genes, including *nahR_2*, and a ribosomal protein L7/L12-like gene (TOL5_08720) were upregulated, suggesting that severe desiccation promotes responses to nucleic acid damage together with adjustments in the translation machinery. In contrast, *cca*, which is involved in tRNA maturation, and the translation-related gene *yegQ* were downregulated, along with a superfamily I DNA/RNA helicase (TOL5_24570) and an XRE-family transcriptional regulator (TOL5_23610). At 52% RH, genes involved in RNA processing, including the nucleotide excision repair factor *uvrB*, the sulfurtransferase *tusA* involved in tRNA 2-thiolation, and multiple translation-related genes (e.g., *ykgM*, *der*, *rpsF*, and *rplI*) were upregulated together with transcriptional regulators (e.g., *nahR_2* and *frmR*). Conversely, several genes involved in replication and translation, including *dnaG*, *yheS*, and *pheT*, as well as *cca* and *yegQ*, were downregulated, indicating concurrent induction and repression within information-processing modules during desiccation. Overall, these DEGs indicate a humidity-dependent shift, with 52% RH associated with broader RNA and translation remodeling alongside genome maintenance, whereas 8% RH is marked by a more damage-focused DNA and RNA repair profile.

DEGs assigned to the “Cellular processes and signaling” supercategory are summarized in Table S3. At 8% RH, genes involved in outer-membrane lipoprotein trafficking (*lolA* and *lolB*) and the protein export chaperone *secB* were upregulated, and increased expression of *dsbC_3*, which is involved in disulfide-bond rearrangement, was also observed. These changes suggest strengthened trafficking and maturation of envelope-associated proteins under severe desiccation. In contrast, *eptA*, which encodes a lipid A phosphoethanolamine transferase, was downregulated, together with signal peptidase I (*lepB*), and an AcrB-family efflux component (*acrB_2*), suggesting selective downshifts in specific envelope functions. At 52% RH, in addition to *lolA* and *secB*, envelope- and stress-related genes, including *tpgA*, *lpxC*, *clpB*, *ahpC_2*, and the betaine–choline–carnitine transporter *betT_2* (38), were also upregulated. In parallel, a type IV pili-related gene *pilV_2* and a FilA adhesin-related gene (TOL5_07310) were also downregulated, suggesting that desiccation drives remodeling of cell-surface structures and outer-membrane composition.

DEGs assigned to the “Metabolism” supercategory are summarized in Table S4. Under both desiccation conditions, Tol 5 exhibited a significant transcriptional downshift in metabolic functions. The most pronounced and consistent decreases were observed in lipid-related functions, particularly fatty acid biosynthesis and acyl-CoA turnover: acetyl-CoA carboxylase components (*accB* and *accD*), fatty acid elongation enzymes (*fabZ* and *fabI*), propionate-CoA ligase (*prpE*), and multiple acyl-CoA dehydrogenases (including TOL5_22700, TOL5_21170, and *ydiO_3*) were downregulated. In parallel, genes linked to aromatic compound uptake and catabolic entry were broadly downregulated, including *benA* and *benK*, as well as the MFS transporter *mucK*, along with additional aromatic-funneling genes (*paaF_2*, *paaH*, *paaJ_2*, and *hpaE*). Furthermore, downregulation of proline utilization genes (*putA* and *putP_1*) and the biotin synthase gene *bioB* aligns with the interpretation that growth- and assimilation-associated metabolism is generally suppressed during desiccation. In contrast, a subset of metabolic genes was upregulated. Under both 8% RH and 52% RH, genes involved in nucleotide-related metabolism (e.g., *prs_2* and *xpt*), a cofactor-related gene encoding a YggS family enzyme (TOL5_09780), and a flavohemoprotein (TOL5_37890) potentially related to redox and gas-responsive functions were upregulated. Furthermore, at 52% RH, upregulation of transport systems involved in nutrient and inorganic ions (e.g., *tauA*, *aroP_2*, *zitB_1* and *ybaL*) was also observed, suggesting that, in parallel with overall metabolic suppression, specific metabolic modules supporting resource acquisition and stress adaptation are selectively activated.

DEGs classified as “Poorly characterized” are summarized in Table S5. At 8% RH, a putative DNA modification/repair radical SAM protein (TOL5_25060) and the ribonucleoprotein *rsr* were upregulated, suggesting that these factors may work together with the DNA/RNA repair pathways described above to support desiccation survival. In addition, a substantial number of COG-unassigned DEGs were detected (23 genes at 8% RH and 31 genes at 52% RH), and most of them were annotated as hypothetical proteins (Table S6). Some of these genes exhibited strong upregulation or downregulation, indicating that Tol 5 desiccation responses include components that are not fully captured by current functional annotations. Accordingly, the poorly characterized and COG-unassigned gene sets represent important targets for expanding the molecular understanding of desiccation tolerance in *Acinetobacter* strains.

## Discussion

In this study, we evaluated the desiccation tolerance of *Acinetobacter* sp. Tol 5, a promising chassis candidate for biomanufacturing, across environments with different humidities, in comparison with the model strains *E. coli* and *P. putida*. At 52% and 8% RH, Tol 5 maintained substantially higher CFU than *E. coli* and *P. putida* (Fig. 1). Consistently, intracellular ATP levels in *E. coli* dropped below the detection limit at 8% and 52% RH, whereas Tol 5, together with *P. putida*, retained comparatively high ATP levels (Fig. 2). In toluene degradation assays, Tol 5 retained toluene-degrading activity after desiccation exposure, with activity slightly higher than that of *P. putida* mt-2 (Fig. 3). In addition, transcriptome profiling revealed that Tol 5 desiccation tolerance is supported not by a single stress-response pathway but by a broad set of mechanisms, including DNA and RNA repair and quality control, oxidative stress regulation, remodeling of the cell envelope, and contributions from poorly characterized proteins (Fig. 4). Together, these results provide both functional and mechanistic evidence that Tol 5 has a high tolerance to desiccation, highlighting its potential as a robust chassis for gas-phase and low-water-content bioprocesses.

Transcriptome profiling indicates that nucleic acid maintenance is a major contributor to Tol 5 desiccation tolerance. Desiccation and subsequent rehydration are known to impose DNA lesions in diverse bacteria, and survival often depends on active repair and recovery programs (31, 39, 40). Consistent with this, Tol 5 exhibited humidity-dependent remodeling within the “information storage and processing” module (Table S2). At 52% RH, the DEG pattern indicates a coordinated maintenance mode that couples genome integrity functions with the tuning of RNA processing and translation capacity, compatible with preserving recovery potential while limiting growth-associated activities. At 8% RH, the response shifts toward a higher-priority nucleic acid protection response, including strong induction of *recA* and the RNA repair ligase *rtcB_2* together with signatures consistent with RNA surveillance (*rsr*) and additional protective factors (Table S2; Table S5). Collectively, these profiles support a staged strategy in which Tol 5 maintains broad maintenance functions under moderate desiccation but reallocates resources toward immediate nucleic acid safeguarding under severe desiccation.

Desiccation is widely associated with enhanced reactive oxygen species formation and oxidative damage, and oxidative stress resistance is intimately tied to survival in classical extremotolerant models (41). However, canonical oxidative-stress response genes that are frequently reported to be induced during desiccation, such as major catalases and other core oxidative-stress defense genes, did not emerge as prominent DEGs under our cutoff criteria. Instead, we observed a consistent induction of a flavohemoprotein gene (TOL5_37890) under both desiccation conditions. Flavohemoproteins are best known as microbial NO-detoxifying enzymes (42, 43) and can influence respiratory redox balance. In addition, downregulation of proline catabolism (*putA*) may favor proline accumulation, and proline has been reported to contribute to tolerance for oxidative and osmotic stress as a compatible solute (44). These suggest that Tol 5 may rely on a less canonical, gas- and redox-responsive strategy during desiccation rather than a broad transcriptional upshift of canonical antioxidant enzymes.

The cell envelope appeared to be actively remodeled under desiccation, as suggested by the induction of outer-membrane lipoprotein trafficking components (*lolA* and *lolB* at 8% RH) and protein export factors such as *secB*, together with increased expression of *dsbC_3* under severe desiccation. The lipoprotein localization (Lol) pathway is essential for outer membrane lipoprotein trafficking and, therefore, may support envelope maintenance under stress (45). In addition, transcripts linked to surface appendages and adhesins changed under desiccation, implying that Tol 5 may adjust cell surface architecture in ways that could influence aggregation, attachment, and microenvironmental hydration. Biofilm and extracellular polymeric substances can retain water and protect cells from desiccation (46), providing a plausible ecological rationale for regulating adhesion-related genes during desiccation.

These transcriptome results suggest that Tol 5 may retain ATP under low humidity primarily by reducing ATP demand and stabilizing membrane energetics, rather than through transcriptional upregulation of ATP generating pathways. First, the broad downregulation of metabolic genes, including those involved in lipid metabolism and aromatic uptake and catabolic entry, indicates a shift away from growth- and assimilation-associated activities that would consume ATP. In parallel, the induction of envelope proteostasis and trafficking factors (e.g., *lolA*, *lolB*, *secB*, *dsbC_3*, and *clpB*) suggest reinforced maturation and quality control of envelope proteins, which could help maintain membrane integrity and limit ion or proton leakage during dehydration. Finally, the upregulation of *ahpC_2* at 52% RH together with a flavohemoprotein (TOL5_37890) points to redox adjustments that could protect respiratory and membrane associated components from oxidative or nitrosative damage, thereby indirectly helping sustain membrane potential and proton motive force. Although these links remain inferential, they offer a plausible explanation for why Tol 5 retains measurable ATP under low humidity conditions despite an overall transcriptional downshift in central metabolic functions.

Taken together, our results not only describe Tol 5’s desiccation response but also nominate a set of engineering targets and design principles for building more robust gas-phase biocatalysts. These include staged nucleic acid maintenance (DNA repair plus RNA repair and surveillance), envelope proteostasis (lipoprotein trafficking, periplasmic maturation, and porin remodeling), redox tuning, and transport rebalancing toward detoxification and homeostasis. Collectively, these elements offer a rational basis for strain improvement and process optimization aimed at sustaining both viability and catalytic performance under low-water-activity conditions.

## Conclusion

In this study, we demonstrated that the toluene-degrading bacterium Tol 5 maintains high viability, cellular energy status, and metabolic activity even after incubation under low-humidity conditions, and we further elucidated the underlying gene-expression responses associated with desiccation stress. Given its strong adhesiveness, versatile metabolism of non-sugar substrates, and high desiccation tolerance, Tol 5 is a promising chassis strain for bioprocesses in low-water-activity environments, including gas-phase bioprocesses. In the future, leveraging the molecular mechanisms identified here to further enhance desiccation tolerance is expected to accelerate the development of high-performance bioprocesses for the conversion of volatile organic compounds.

## Declarations

### Ethics approval and consent to participate

Not applicable.

### Consent for publication

Not applicable.

### Availability of data and materials

RNA-seq data reported are available in the DDBJ Sequenced Read Archive under the accession numbers PRJDB39957.

### Authors’ contributions

SY, SI, and KH designed the study. HY performed CFU, ATP, and toluene degradation assay. SY, HY, and SI performed transcriptome analysis. SY and HY wrote the draft manuscript and all authors reviewed and approved the final manuscript.

## Acknowledgements

The authors wish to acknowledge the Center for Gene Research, Nagoya University, for technical support with the RNA sequencing.

