## Supplementary Information for "Desiccation-tolerant *Acinetobacter* as a robust chassis for gas-phase bioprocesses"

**Table S1. Gene expression in Tol 5 under different desiccation conditions.**

**Table S2. DEGs assigned to the "Information storage and processing" supercategory.**

**Table S3. DEGs assigned to the "Cellular processes and signaling" supercategory.**

**Table S4. DEGs assigned to the "Metabolism" supercategory.**

**Table S5. DEGs classified as "Poorly characterized."**

**Table S6. DEGs unassigned to any COG category.**
